# Long-Term Functional and Histological Outcomes Following Sutureless Peripheral Nerve Repair Using Nerve Tape in Pigs

**DOI:** 10.64898/2026.07.29.741584

**Authors:** Mykhailo M. Tatarchuk, David R. Clizbe, Kevin D. Browne, Susanna Howard, Yohannes Ghenbot, Eric L. Zager, D. Kacy Cullen, Justin C. Burrell

## Abstract

Severe peripheral nerve injuries result in incomplete recovery despite neurorrhaphy. Microsurgical suturing is technically demanding, time-intensive, and may produce variable fascicular alignment. Nerve Tape is an FDA-approved sutureless device enabling rapid, reproducible nerve coaptation. This study compared Nerve Tape with epineurial microsuturing following common peroneal nerve transection in Yucatan minipigs. Over 12 months, both groups demonstrated reinnervation of the tibialis anterior and extensor digitorum brevis, representing proximal and distal muscle targets, respectively. Tibialis anterior recovery was comparable between groups. In contrast, Nerve Tape produced greater distal motor recovery in the extensor digitorum brevis, with approximately 1.8-fold higher compound muscle action potential amplitude and 74.3% versus 46.0% recovery compared with microsutures. Compound nerve action potential amplitudes recorded from the motor branch of the deep peroneal nerve were also greater with Nerve Tape, whereas conduction velocities were comparable. Histological analysis demonstrated preserved fascicular architecture distal to the repair in both groups, with no significant differences in axon count, mean myelinated axon diameter, or g-ratio in the terminal common peroneal nerve or its distal motor branch. Clinical use was demonstrated in a representative case with progressive recovery. Nerve Tape supported durable structural and functional recovery and improved distal motor reinnervation compared with microsuturing.

## Introduction

Peripheral nerve injuries (PNIs) affect millions of individuals worldwide each year and frequently result in persistent motor deficits despite surgical intervention.^1^ The current standard of care, microsurgical epineurial suturing, is technically demanding, time intensive, and highly dependent on operator skill.^2^ Even under optimal conditions, achieving precise fascicular alignment remains challenging, and variability in coaptation quality contributes to inconsistent functional outcomes.^3^ In addition, surgical manipulation and suturing at the repair site is associated with aberrant inflammatory and fibrotic responses which may impede nerve healing and regeneration.

To address these limitations, numerous sutureless and adjunctive repair strategies have been explored, including fibrin sealants, laser-assisted coaptation, nerve wraps, and bioadhesive materials.^4^ While many of these approaches have demonstrated feasibility in small animal models, few have shown robust distal regeneration or durable long-term functional outcomes in large animal systems that more closely approximate human nerve size and regenerative kinetics.

Nerve Tape^®^ is an FDA-approved, commercially available repair device composed of flexible columns of Nitinol microhooks embedded between laminated sheets of processed porcine small intestine submucosa (SIS).^5^ For use in the coaptation of severed nerve, the nerve stumps are aligned by positioning on opposing microhooks, which engage the outer epineurium to provide immediate stabilization. The device is then wrapped circumferentially around the coaptation site, enabling additional microhook engagement and structural reinforcement while creating a contained repair environment. The SIS component serves as a biologically-derived scaffold that supports cellular infiltration and tissue remodeling.^6^ Prior studies in small animal models have demonstrated favorable handling characteristics and biomechanical strength relative to sutured repair^7^, and cadaveric studies have highlighted improvements in procedural efficiency.^8^ However, the long-term regenerative performance of this approach in a translational large animal model has not been comprehensively evaluated.

In this study, we assessed 12-month electrophysiological and histological outcomes following Nerve Tape-mediated repair of the common peroneal nerve (CPN) in Yucatan minipigs, a large animal model previously established and validated by our group to approximate human peripheral nerve caliber and regenerative dynamics.^9^ Here, we apply this model with long recovery time points and the incorporation of multimodal outcome measures, including electrophysiology, distal target reinnervation, and histological assessment of regeneration and tissue response. We anticipated that Nerve Tape would support functional and structural recovery comparable to that achieved with standard neurorrhaphy featuring epineurial suturing.

## Methods

The animal study was approved by the University of Pennsylvania Institutional Animal Care and Use Committee (IACUC). The clinical vignette was conducted in accordance with protocols approved by the University of Pennsylvania Institutional Review Board (IRB), and written informed consent was obtained from the patient for publication of clinical details and associated media.

Ten female adolescent Yucatan minipigs (6–8 weeks old, 30–35 kg) were enrolled in a randomized controlled study evaluating long-term functional and histological outcomes following CPN transection and immediate repair. Animals were randomly assigned to Nerve Tape repair (n = 5) or standard epineurial repair using 8–10 interrupted 8-0 prolene sutures (n = 5).

### Surgical Approach

All procedures were performed under general anesthesia using aseptic technique. Animals were anesthetized with ketamine (20 mg/kg, intramuscular) and midazolam (0.4 mg/kg, intramuscular). Anesthesia was induced and maintained with isoflurane (5% induction, 1–2% maintenance) followed by endotracheal intubation. Local analgesia was provided with bupivacaine (1–2 mg/kg) infiltrated along the incision. Intraoperative analgesia included meloxicam (0.4 mg/kg, intramuscular) and extended-release buprenorphine (0.1 mg/kg, subcutaneous) and Maropitant (1 mg/kg, subcutaneous) was administered for antiemetic support. Postoperative analgesia consisted of meloxicam (0.1 mg/kg, oral) once daily for 2 days.

The CPN was exposed through a longitudinal lateral thigh incision centered over the fibular head. Blunt and sharp dissection was carried through subcutaneous tissue and fascia to identify the nerve as it courses around the fibular neck. The nerve was mobilized proximally and distally with care to preserve surrounding vasculature. Following isolation, vessel loops were placed to facilitate atraumatic handling. A complete transection was performed at a standardized location proximal to the fibular head. Repair was performed according to group assignment, ensuring tension-free coaptation with careful alignment of the nerve stumps. The wound was irrigated and closed in layers (3-0 vicryl for fascia and subcutaneous tissue, 2-0 PDS for skin) and dressed with Telfa, topical antibiotic ointment, and Tegaderm prior to recovery.

### Longitudinal Non-Invasive Electrophysiological Assessments

Animals were anesthetized with ketamine (10 mg/kg, intramuscular) and midazolam (0.4 mg/kg, intramuscular) and maintained on isoflurane during recordings. Measurements were obtained 15– 30 minutes following induction.

Motor recovery was assessed non-invasively using compound muscle action potentials (CMAPs) recorded from the tibialis anterior (TA) and extensor digitorum brevis (EDB) muscles – a proximal and distal muscle target, respectively – at 2, 4, 6, 9, and 12 months postoperatively. Transcutaneous stimulation of the CPN was delivered using a handheld surface probe (Natus Viking EDX system) with a pulse width of 2.0 ms, amplitude of 0–10 mA, and frequency of 1 Hz. Recordings were obtained using subdermal needle electrodes placed in the muscle belly, with a reference electrode positioned in the distal tendon and a ground electrode placed subcutaneously. Stimulus intensity was increased to achieve supramaximal activation. Peak-to-peak amplitudes were quantified and normalized to the contralateral limb to calculate percent recovery at each time point.

### Terminal Intraoperative Electrophysiology

At 12-months, terminal electrophysiological studies were performed under general anesthesia. The CPN and its distal branches, including the motor deep peroneal nerve (mDPN) and sensory deep peroneal nerve (sDPN), were re-exposed.

CMAPs were recorded from TA and EDB muscles during proximal and branch-level stimulation. For compound nerve action potential (CNAP) recordings, the nerve was stimulated 5 mm proximal to the repair site using a bipolar hook electrode (Rochester Electro-Medical; #400900). CNAPs were recorded 5 mm distal to the repair and approximately 1 cm distal from the bifurcation of the motor branch and sensory branch of the CPN (mDPN and sDPN, respectively) using a bipolar electrode configured to maintain consistent nerve contact. Stimulation parameters were biphasic pulses at 0–1 mA, pulse width 0.2 ms, and frequency 1 Hz. Signals were acquired and bandpass filtered from 10–3000 Hz. A ground electrode was placed in subcutaneous tissue between stimulation and recording sites. Trains of five stimuli were averaged to improve signal-to-noise ratio. Peak-to-peak amplitude and latency were quantified. Conduction velocity was calculated as the distance between stimulation and recording sites divided by latency.

### Tissue Harvest and Histological Analysis

At the 12-month endpoint, distal nerve segments were harvested for histological analysis. Samples were fixed in 2.5% glutaraldehyde, post-fixed in osmium tetroxide, embedded in resin, sectioned at 0.5 μm, and stained with toluidine blue. Quantitative morphometric analyses were performed on distal nerve cross-sections using QuPath software on coded specimens by investigators blinded to treatment group.

For automated axon morphometric quantification, a region of interest captured for a target pixel resolution of 1440 wide or higher was selected for each animal and analyzed using an automated segmentation process based on previously validated methods used in AxonDeepSeg^10^ software analysis. Used on this dataset was DeepAxon-Legacy^11^, further developed by researchers at the VCU Orthopedics Microsurgery Lab and is distributed through an MIT Open Source License and has been additionally trained for peripheral nerve in mouse, rat, and rabbit, and is being used, here, for porcine model analysis, specifically axon count, G Ratio, and Axon Diameter for myelinated axons.

## Statistical Analysis

All analyses were performed on coded data with investigators blinded to treatment group. Data are presented as mean ± SEM. Longitudinal CMAP data were analyzed using a mixed-effects model (REML) in GraphPad Prism 8.0 with fixed effects for treatment and time and a treatment × time interaction. In the absence of missing data, this approach is equivalent to a two-way repeated-measures ANOVA. Post hoc multiple comparisons were performed with appropriate correction. Endpoint comparisons were assessed using two-tailed unpaired Student’s t-tests. Statistical significance was defined as p < 0.05.

## Results

### Transcutaneous CMAP Recovery Over Time

Animals undergoing nerve repair using either Nerve Tape or microsutures demonstrated progressive motor recovery in the TA and EDB muscles over 12 months (**Figure 2**). In the TA, recovery kinetics were similar between the repair groups, with no significant differences at any of the timepoints. In particular, at 12 months post-repair, CMAP amplitudes (14.7 ± 1.7 mV vs. 13.34 ± 0.25 mV, p = 0.9748) and percent recovery (76.8 ± 8.2% vs. 69.3 ± 2.0%, p = 0.9762) were comparable between groups.

**Figure 1.**
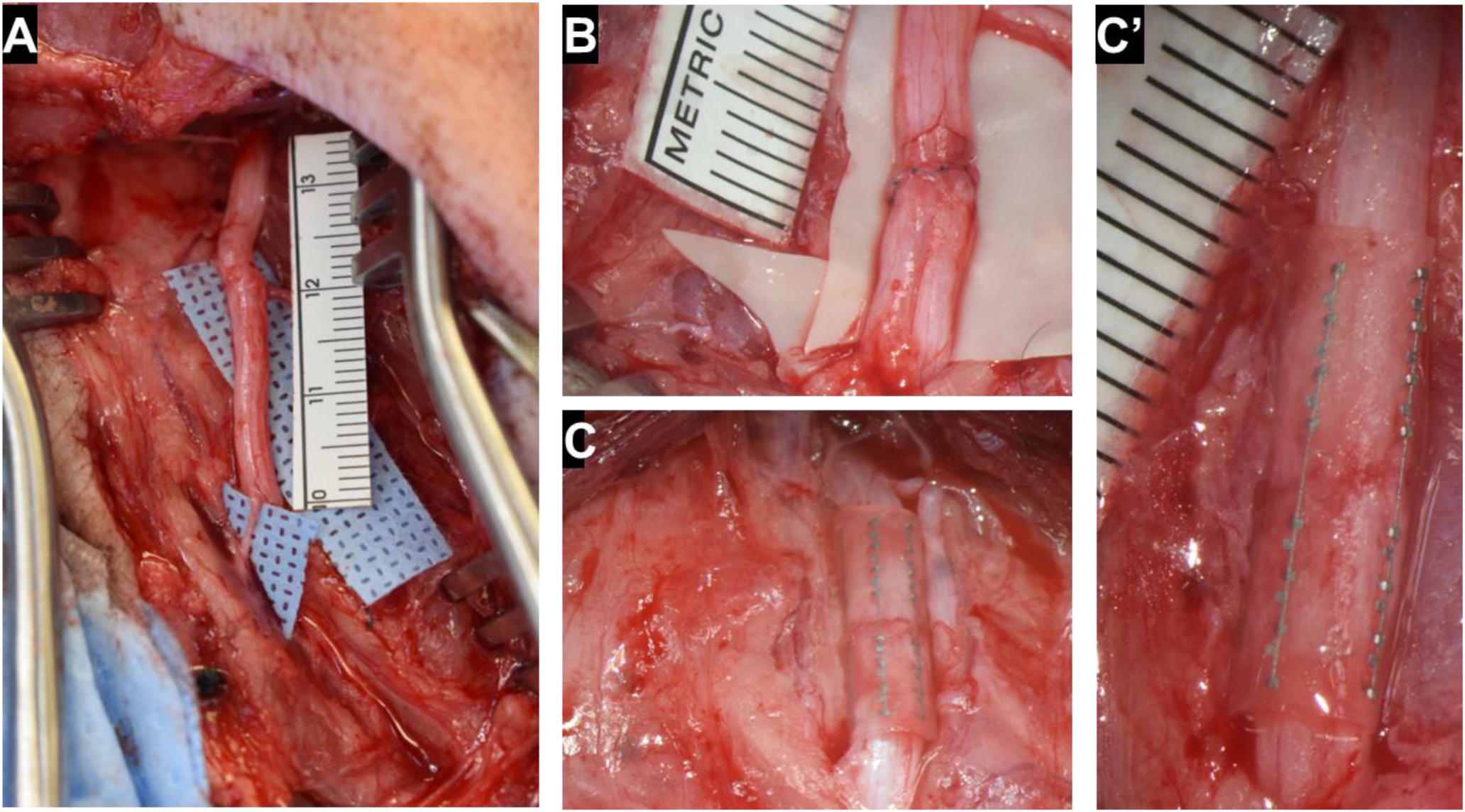
Intraoperative repair of the common peroneal nerve (CPN) using Nerve Tape and epineurial sutures. (A) Intraoperative exposure of the CPN prior to transection. (B) Standard epineurial repair using interrupted 8-0 prolene sutures placed circumferentially at the coaptation interface. (C) Nerve Tape is positioned circumferentially around the coaptation site to enable tension-free alignment, and stable apposition of proximal and distal nerve stumps. (C′) High-magnification view showing Nitinol microhook placement and tissue apposition following Nerve Tape application.

**Figure 2.**
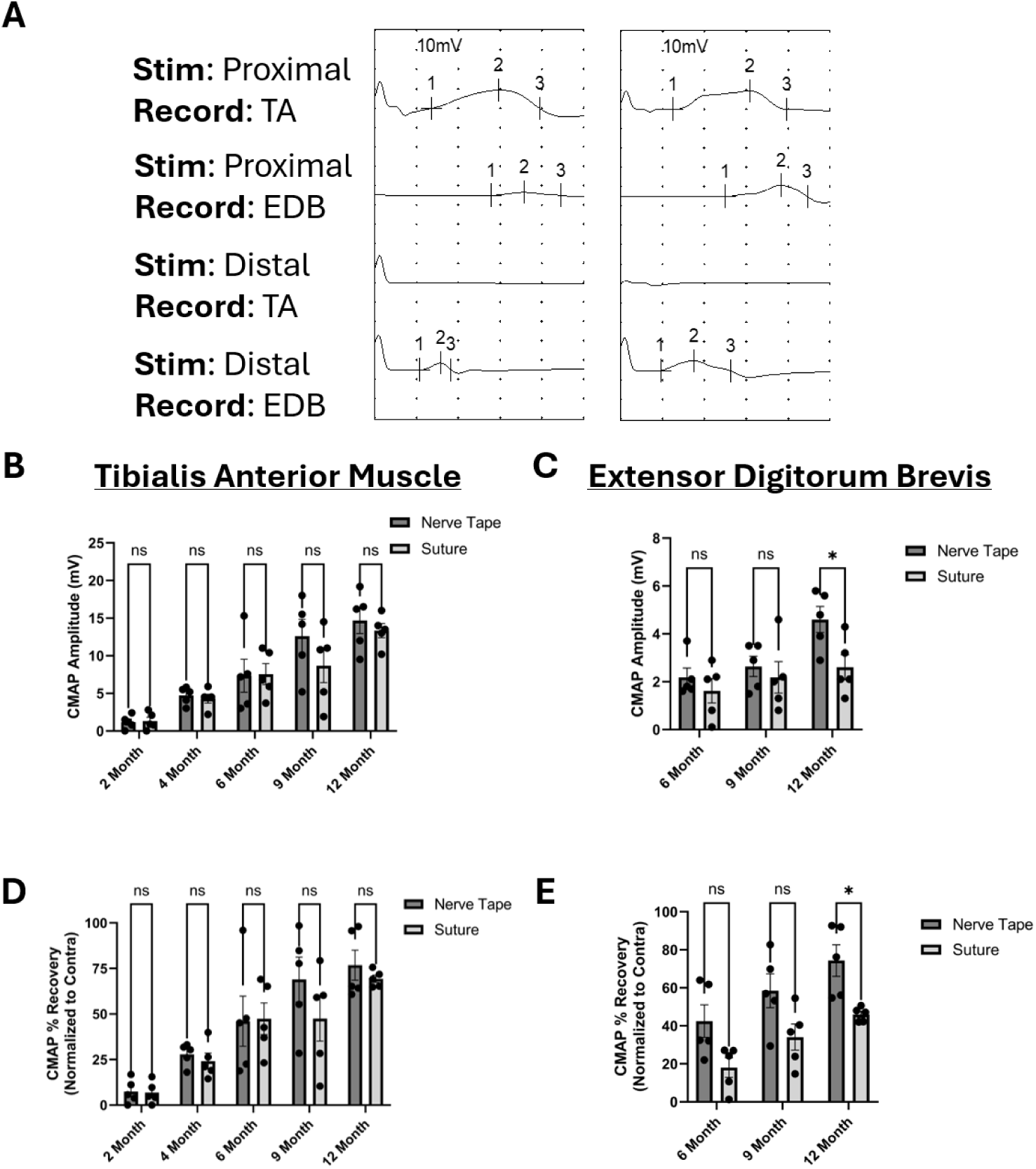
Serial compound muscle action potential (CMAP) recovery in the anterior tibialis (TA) and extensor digitorum brevis (EDB) following common peroneal nerve (CPN) repair. (A) Representative CMAP recordings obtained following proximal CPN stimulation with distal recording from the TA and EDB muscles. (B) TA CMAP amplitudes measured at 2, 4, 6, 9, and 12 months postoperatively demonstrate progressive recovery in both Nerve Tape and suture groups, with no significant differences between groups at any time point. (C) EDB CMAP amplitudes measured at 6, 9, and 12 months demonstrate significantly greater amplitudes in the Nerve Tape group at 12 months (*p* < 0.05). (D) TA CMAP percent recovery normalized to the contralateral limb demonstrates progressive improvement over time, with no significant differences between groups. (E) EDB CMAP percent recovery normalized to the contralateral limb demonstrates significantly greater recovery in the Nerve Tape group at 12 months (*p* < 0.05). Data are presented as mean ± SEM with individual animal values shown. Statistical analyses were performed using two-way repeated-measures ANOVA with Šidák correction for multiple comparisons; “ns” indicates not significant, and * indicates *p* < 0.05.

However, in the EDB muscle, the Nerve Tape group demonstrated significantly greater recovery at 12 months post-repair. Specifically, CMAP amplitude was higher in Nerve Tape animals compared to suture controls (4.60 ± 0.54 mV vs. 2.60 ± 0.52 mV, p = 0.0327). Percent recovery relative to the contralateral side was also greater in the Nerve Tape group (74.3 ± 8.3% vs. 46.0 ± 1.7%, p = 0.0266).

### Gross Morphology at 12 Months

At terminal harvest, Nerve Tape repairs were readily identifiable, with preserved nerve contour and well-defined repair interfaces (**Figure 3**). Qualitative inspection demonstrated minimal surrounding fibrosis, and the device could be dissected from adjacent tissue planes without difficulty. The small intestinal submucosa (SIS) component appeared completely remodeled at 12 months, with no gross evidence of persistent material or encapsulation. Standard epineurial suture repairs demonstrated visible perineural adhesions at the coaptation site and required careful dissection to separate the nerve from surrounding tissue.

**Figure 3.**
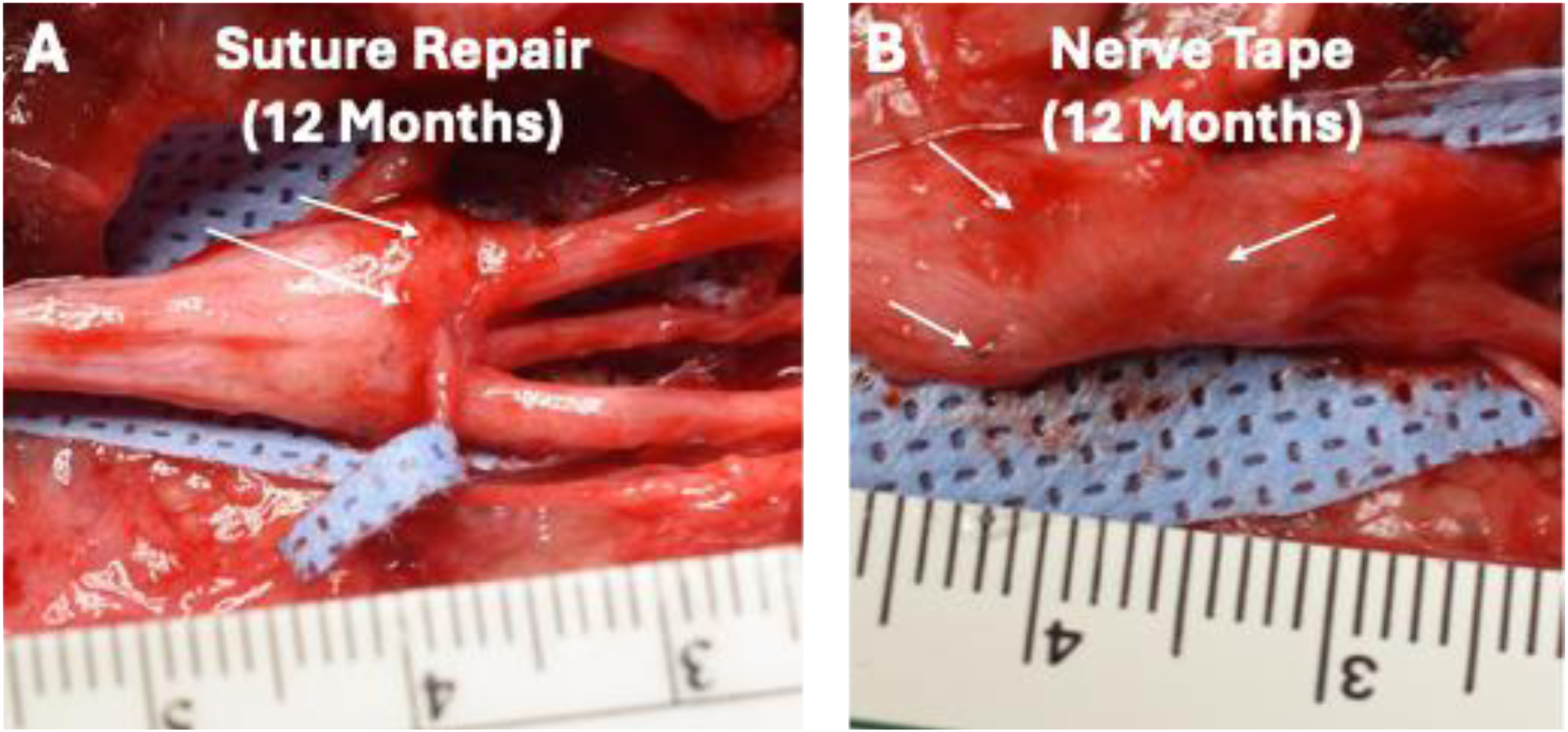
Comparison of coaptation sites at 12 months following suture repair and Nerve Tape repair of the common peroneal nerve (CPN). (A) Standard epineurial suture repair showing irregular remodeling at the coaptation site with visible perineural adhesions and tethering to the surrounding soft tissue, indicated by arrows. (B) Nerve Tape repair showing a preserved nerve contour, a smoother repair interface, and limited gross fibrotic attachment to the surrounding tissue. Arrows indicate the locations of Nerve Tape microhooks incorporated within the remodeled repair site. Compared with the representative sutured repair, the Nerve Tape repair exhibited less apparent perineural adhesion formation and more uniform structural continuity.

### Intraoperative CMAPs at 12 Months

At the 12-month terminal assessment, intraoperative CMAPs from TA following CPN stimulation were similar between groups (**Figure 4**): 7.25 ± 1.06 mV (Nerve Tape) vs. 8.24 ± 2.54 mV (Suture), (*p* = 0.7534). In contrast, EDB responses were significantly higher in the Nerve Tape group for both CPN stimulation (4.53 ± 0.69 mV vs. 2.12 ± 0.29 mV, *p* = 0.0103) and mDPN stimulation (4.48 ± 0.81 mV vs. 2.12 ± 0.55 mV, *p* = 0.0421).

**Figure 4.**
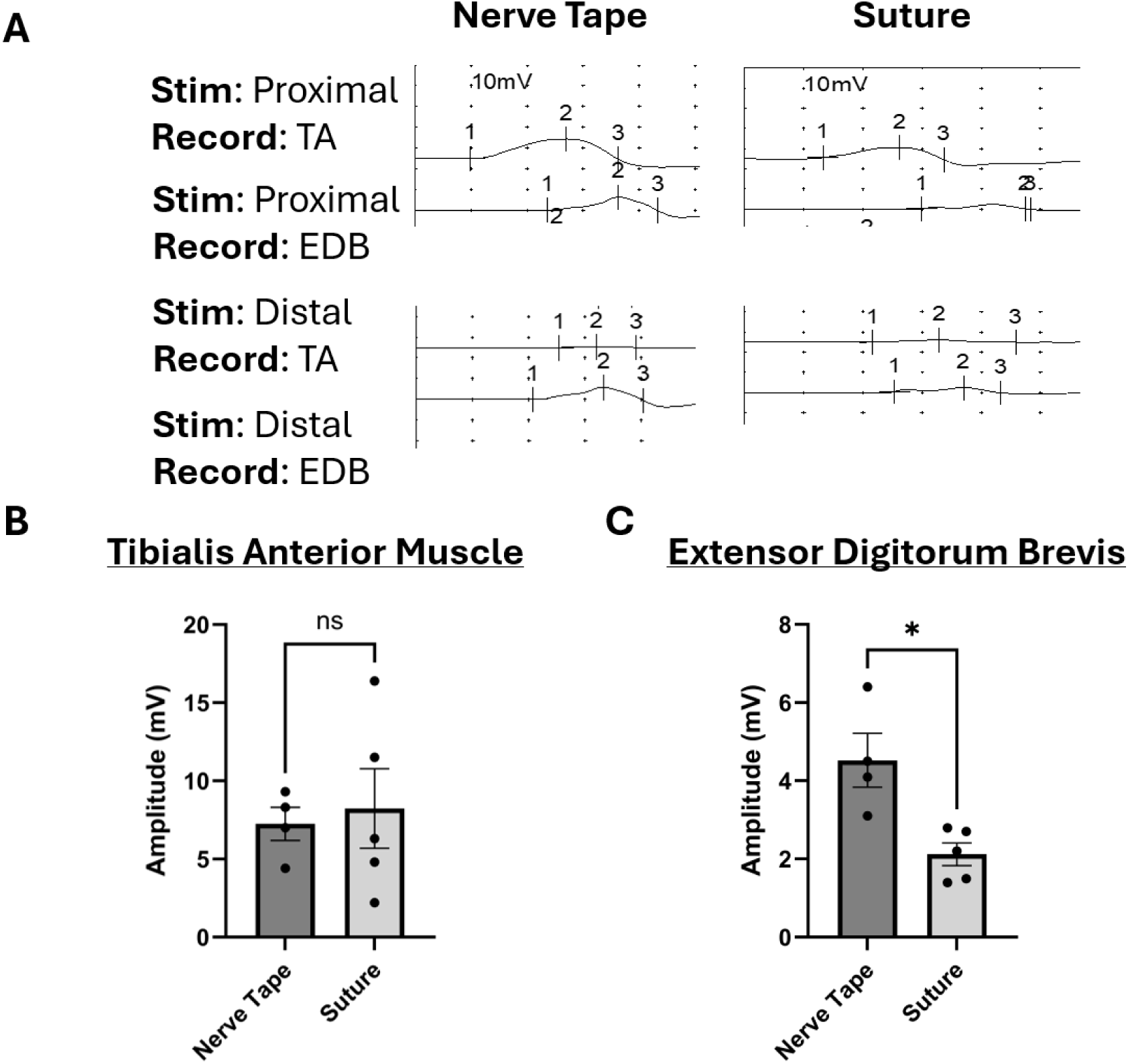
Intraoperative compound muscle action potential at 12 months following common peroneal nerve repair. (A) Representative intraoperative CMAP recordings obtained at 12 months following stimulation of the proximal or distal common peroneal nerve with recordings from the tibialis anterior (TA) and extensor digitorum brevis (EDB) muscles in nerve tape and suture repair groups. (B) TA CMAP amplitudes measured intraoperatively at 12 months demonstrate comparable recovery between nerve tape and suture groups, with no significant difference between groups. (C) EDB CMAP amplitudes measured intraoperatively at 12 months demonstrate significantly greater amplitudes in the nerve tape group compared with the suture group (*p* < 0.05). Data are presented as mean ± SEM with individual data points shown. Statistical comparisons were performed using an unpaired two-tailed *t* test; “ns” indicates not significant, and * indicates *p* < 0.05.

### Intraoperative CNAPs and Conduction Velocity at 12 Months

At 12 months post repair, the CNAP amplitude measured from the CPN trunk were not significantly different between groups (**Figure 5**). However, Nerve Tape repairs showed significantly greater CNAP amplitudes at both motor and sensory branches (mDPN and sDPN, respectively) compared to Suture repairs. From CPN to mDPN, amplitudes were 531 ± 132 µV vs. 187.5 ± 25.7 µV (*p* = 0.0339), and from CPN to sDPN, 423.1 ± 35.5 µV vs. 279.5 ± 21.2 µV (*p* = 0.0084). Conduction velocities were similar between groups. From CPN to mDPN, Nerve Tape and Suture repairs yielded velocities of 52.4 ± 2.2 m/s and 51.25 ± 3.4 m/s, respectively (*p* = 0.7782). Sensory conduction velocity from CPN to sDPN was also comparable (47.2 ± 0.8 m/s vs. 49.5 ± 3.2 m/s, *p* = 0.4586), suggesting similar speed of propagation across the repair. Higher signal amplitudes in the Nerve Tape group with comparable conduction velocities suggest improved axonal conduction and maturation.

**Figure 5.**
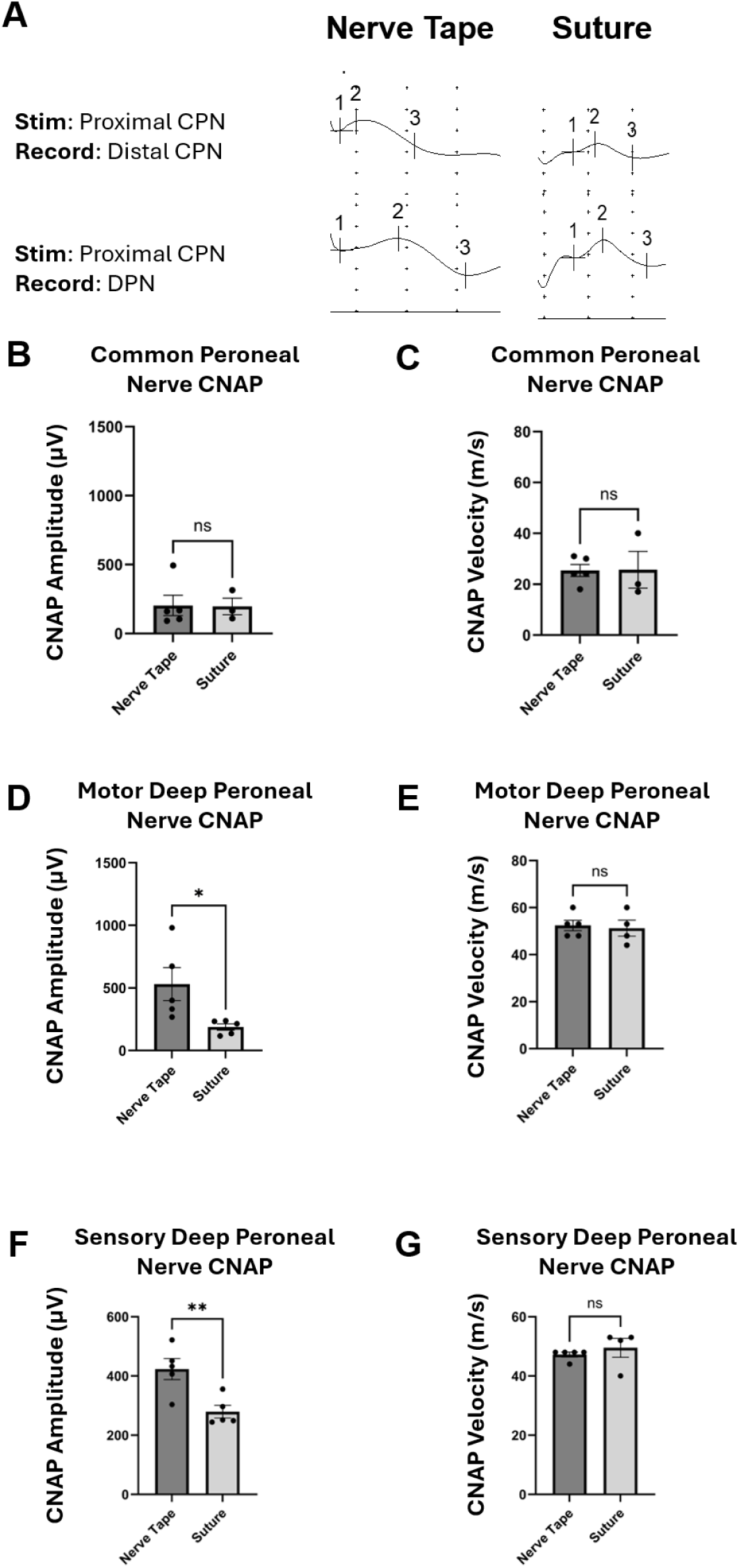
Terminal intraoperative compound nerve action potential (CNAP) amplitude and conduction velocity at 12 months. (A) Representative CNAP waveforms recorded from the terminal branch of the common peroneal nerve (CPN; top) and the distal motor branch of the deep peroneal nerve (mDPN; bottom). (B) CNAP amplitudes recorded from the terminal CPN branch distal to the repair site demonstrate significantly greater amplitudes in the nerve tape group compared with the suture group (*p* < 0.05). (C) Conduction velocities calculated from proximal stimulation to the terminal CPN branch were comparable between groups. (D) CNAP amplitudes recorded from the distal motor branch of the deep peroneal nerve demonstrate significantly greater amplitudes in the nerve tape group compared with the suture group (*p* < 0.05). (E) Conduction velocities calculated from proximal stimulation to the distal mDPN were comparable between groups. (F) CNAP amplitudes recorded from the distal sensory branch of the deep peroneal nerve demonstrate significantly greater amplitudes in the nerve tape group compared with the suture group (*p* < 0.05). (G) Conduction velocities calculated from proximal stimulation to the distal sensory branch of the deep peroneal nerve were comparable between groups. Data are presented as mean ± SEM with individual animal values shown. Statistical comparisons were performed using unpaired two-tailed *t* tests; “ns” indicates not significant, * indicates *p* < 0.05, and ** indicates *p* < 0.01. Greater distal CNAP amplitudes in nerve tape-treated animals, without corresponding differences in conduction velocity, are consistent with a greater number of conducting axons reaching distal motor and sensory branches rather than altered intrinsic conduction properties.

### Histological Outcomes at 12 Months

Toluidine blue–stained semithin nerve sections demonstrated preserved fascicular architecture distal to the repair in both groups.

Automated quantitative morphometry analysis of the terminal branch of the common peroneal nerve (CPN) demonstrated no significant differences between groups (**Figure 6**). Axon counts were not statistically different, but showed a strong trend towards increased axon counts following suture repair (suture: 666.3 ± 128.2 vs Nerve Tape: 295 ± 76.97, p =0.0594). Myelinated axon diameter (2.615 ± 0.1907 vs 3.239 ± 0.3981, p =0.2543), and g-ratio (0.5418 ± 0.01974 vs 0.5603 ± 0.02099, p =0.5504) demonstrated comparable axonal caliber and myelination profiles within the CPN.

**Figure 6.**
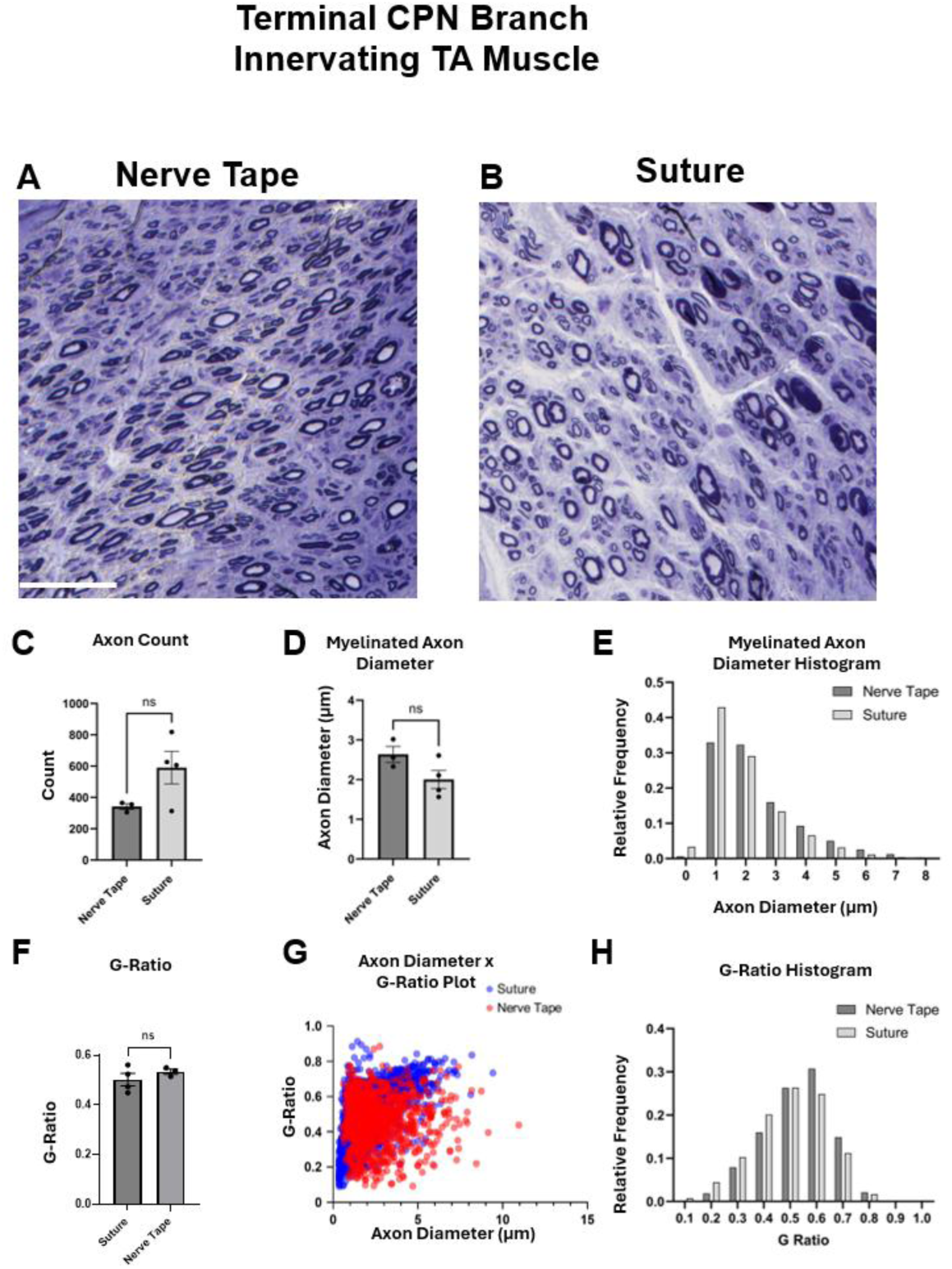
Histological assessment of the terminal branch of the CPN at 12 months. (A) Representative toluidine blue-stained semithin section of the terminal branch of the common peroneal nerve (CPN), distal to the repair site, following Nerve Tape repair. Scale bar shown at 20 um. (B) Representative toluidine blue-stained semithin section of the corresponding terminal CPN branch following conventional epineurial suture repair. Both groups demonstrate preserved fascicular architecture and heterogeneous populations of myelinated axons. (C) Quantification of axon counts in the terminal CPN branch demonstrating no significant difference between groups. (D) Mean myelinated axon diameter in the terminal CPN branch, showing no significant difference between repair groups. (E) Frequency distribution of myelinated axon diameters in the terminal CPN branch. (F) Mean g-ratio in the terminal CPN branch, demonstrating no significant difference between groups. (G) Scatter plot of myelinated fiber diameter versus g-ratio for individual fibers. (H) Frequency distribution of g-ratio values in the terminal CPN branch. Across all measured morphometric parameters, including axon counts, myelinated axon diameter, and g-ratio, no significant differences were observed between Nerve Tape and suture repairs. Data are presented as mean ± SEM with individual animal values shown. Statistical comparisons were performed using unpaired two-tailed *t* tests; “ns” indicates not significant.

Similarly, the automated segmentation analysis of the mDPN (distal branch of the CPN) demonstrated no significant differences between Suture vs Nerve Tape (**Figure 7**). The axon count (Suture: 600.2 ± 129.8 vs 778.8 ± 172.5, p=0.4339), mean myelinated axon diameter (2.264 ± 0.318 vs 1.956 ± 0.1578, p=0.4194), and g-ratios (0.4867 ± 0.0304 vs 0.4982 ± 0.0078, p=0.7297) showed similar profiles as well.

**Figure 7.**
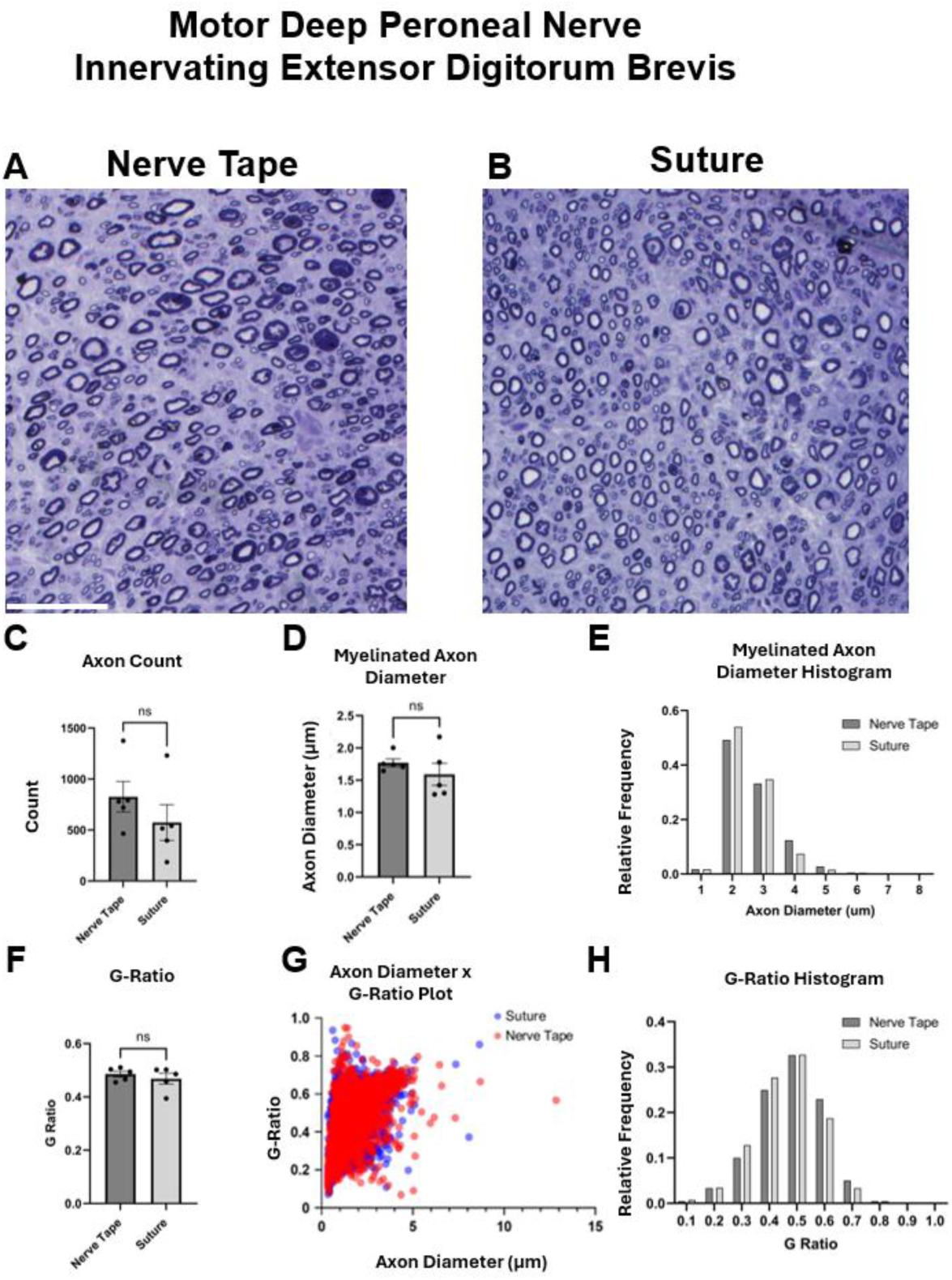
Histological assessment of the distal motor branch of the deep peroneal nerve at 12 months following common peroneal nerve repair. (A) Representative toluidine blue-stained semithin section of the distal motor branch of the deep peroneal nerve (mDPN) following Nerve Tape repair. Scale bar shown at 20 um. (B) Representative toluidine blue-stained semithin section of the distal mDPN following conventional epineurial suture repair. Both groups demonstrate preserved fascicular architecture and heterogeneous populations of myelinated axons. (C) Quantification of axon counts in the distal mDPN demonstrating no significant difference between groups. (D) Mean myelinated axon diameter in the distal mDPN demonstrating no significant difference between groups. (E) Frequency distribution of myelinated axon diameters in the distal mDPN. (F) Mean g-ratio in the distal mDPN demonstrating no significant difference between groups. (G) Scatter plot of myelinated fiber diameter versus g-ratio for individual fibers. (H) Frequency distribution of g-ratio values in the distal mDPN demonstrating comparable myelination profiles between groups. Across all measured morphometric parameters, including axon counts, myelinated axon diameter, and g-ratio, no significant differences were observed between Nerve Tape and suture repairs. Data are presented as mean ± SEM with individual animal values shown. Statistical comparisons were performed using unpaired two-tailed *t* tests; “ns” indicates not significant.

### Clinical Vignette

To demonstrate the clinical use of Nerve Tape and to contextualize the importance of distal reinnervation, we present a representative clinical vignette. A 72-year-old male presented with five months of progressive left upper extremity weakness following a ground-level fall. Prior C4– 6 anterior cervical discectomy and fusion did not improve symptoms. Examination demonstrated severe proximal weakness (MRC scale: deltoid 0/5, biceps 0/5) with preserved distal function. Electromyography confirmed chronic C5–6 radiculopathy with active denervation and no electrical continuity to the deltoid or biceps.

The patient underwent staged nerve transfer procedures, including an Oberlin transfer^12^ (internal neurolysis of the musculocutaneous and ulnar nerves with fascicular transfer) followed by a radial nerve triceps branch transfer to the axillary nerve.^13^ Nerve coaptation in these procedures was performed using Nerve Tape as a sutureless repair interface (**Figure 8; Supplementary Video 1**). Postoperatively, the patient experienced transient radial forearm pain that resolved spontaneously.

**Figure 8.**
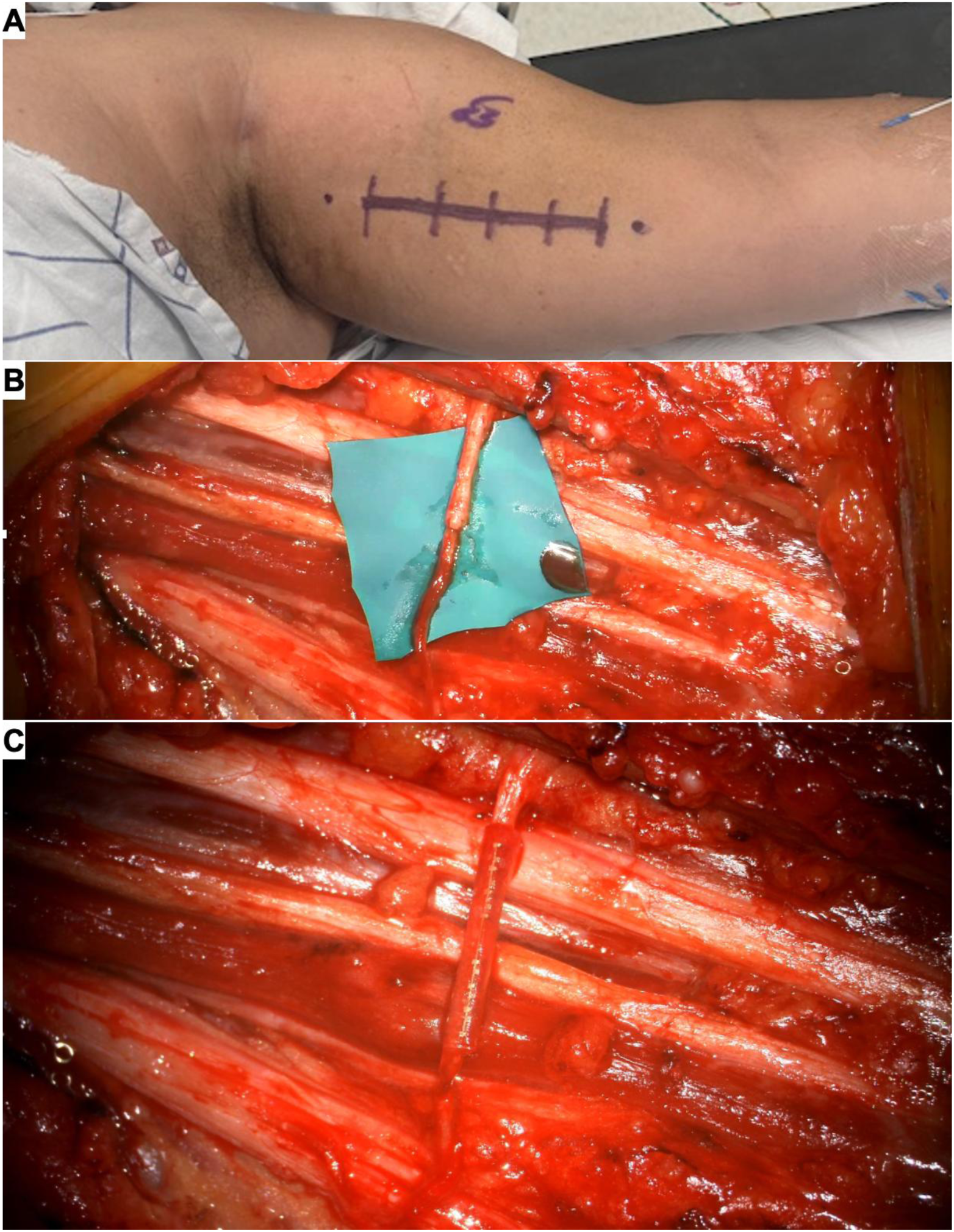
Sutureless Oberlin nerve transfer using Nerve Tape in a clinical nerve transfer. (B) Preoperative surface marking of the surgical site along the proximal upper arm. (C) Intraoperative exposure of the donor ulnar nerve fascicle and recipient motor branch to the biceps following microsurgical dissection. A protective background was positioned beneath the nerves before coaptation. (D) Completed sutureless Oberlin transfer using Nerve Tape, securing the donor ulnar nerve fascicle to the motor branch of the biceps without epineurial sutures while maintaining alignment and stabilization across the coaptation interface.

Over the subsequent year, the patient recovered antigravity function in both the deltoid and biceps without loss of donor nerve function, with preserved strength in the triceps, brachioradialis, and hand intrinsics. This case shows the straightforward and effective use of Nerve Tape in a complex nerve transfer procedure, while highlighting the importance of successful distal motor reinnervation in functional recovery following nerve injury.

## Discussion

Sutureless strategies for peripheral nerve repair have been explored for decades, including fibrin sealants, laser-assisted coaptation, bioadhesive hydrogels, and nerve wraps.^4, 14–16^ While many have shown feasibility in rodent models and short-term studies, durable large-animal data with long-term functional and structural endpoints remain limited.

In this study, a Yucatan minipig common peroneal nerve repair model, previously developed by our group^9, 17^, was used as a translationally relevant preclinical platform. This model enables evaluation of long regeneration distances, delayed functional recovery, and longitudinal electrophysiological outcomes – hallmark features of peripheral nerve regeneration that are not typically replicated in small animal models. Indeed, the porcine CPN shares key anatomical and physiological features with human mixed motor nerves, while permitting extended follow-up and serial assessments. In applying this porcine model to evaluate the use and efficacy of Nerve Tape for direct coaptation, we found that Nerve Tape supported robust axonal regeneration and functional recovery, with electrophysiological and histomorphometry findings consistent across proximal and distal targets.

The overall goal of this study was to compare conventional microsuture neurorrhaphy with Nerve Tape as a sutureless alternative, with the expectation that Nerve Tape would demonstrate non-inferiority, or efficacy comparable to standard repair. However, animals undergoing primary nerve repair with Nerve Tape demonstrated superior outcomes across several key electrophysiological measures. These differences were most pronounced at distal targets, where Nerve Tape repair produced greater CMAP amplitudes and percent recovery in the EDB, together with greater CNAP amplitudes recorded from the distal motor and sensory branches of the deep peroneal nerve. In contrast, axon counts, mean myelinated axon diameter, g-ratio, and conduction velocity were comparable between repair groups. There was also a strong, but non-significant, trend toward greater axon counts in the terminal CPN following suture repair. Although greater axon counts in the terminal CPN might initially appear to indicate enhanced regeneration, elevated fiber counts distal to a repair can also reflect collateral sprouting and incomplete pruning of supernumerary regenerative branches rather than a greater number of parent axons successfully reinnervating distal targets.^18–20^

The combination of greater distal electrophysiological amplitudes with comparable conduction velocity and histomorphometry suggests that Nerve Tape may improve the functional yield of regeneration without adversely affecting axonal maturation or myelination. Because compound response amplitude is influenced by the number of activated fibers, their physiological properties, and the temporal synchrony of their action potentials, these findings are consistent with more effective functional continuity and distal target reinnervation.^21, 22^ Given that prolonged regeneration distances and inaccurate target reinnervation are major limitations to functional recovery after peripheral nerve injury, these results suggest that the quality of the repair interface may influence how effectively regenerating axons establish functional connections with terminal motor and sensory targets.^23, 24^

The distal specificity of these effects may reflect the inherent demands of this large-animal model of peripheral nerve injury. Longer regeneration distances create a more stringent setting in which subtle differences in repair quality may become apparent. When repairs are performed by an experienced surgeon, proximal outcomes may converge, whereas differences may emerge only after axons must regenerate over extended distances to reach terminal targets. Prior work has also suggested that Nerve Tape may improve coaptation alignment and reduce technical variability. Although these features were not directly assessed in the present study, they may contribute to the improved distal electrophysiological recovery observed here.

Device-related tissue response was not a primary endpoint of the present study. Although Nerve Tape introduces additional biomaterial at the coaptation site, its components have demonstrated favorable remodeling characteristics in prior preclinical studies, including limited fibrosis and macrophage accumulation at the repair interface.^25^ Although operative time was not prospectively recorded in the present study, previous investigations have consistently demonstrated that, after surgeons become familiar with the device, Nerve Tape substantially reduces nerve coaptation time compared with conventional microsuture repair. In a recent clinical evaluation involving attending surgeons and trainees, mean coaptation time was reduced from 5.20 to 1.79 minutes, representing an approximately 66% reduction while simultaneously improving the rate of clinically acceptable repairs.^26^ Given that peripheral nerve reconstruction often requires multiple coaptations, reductions in repair time of this magnitude have the potential to meaningfully decrease operating room utilization and associated procedural costs. Consequently, decreased operative time may partially offset the increased device cost of Nerve Tape relative to microsuture repair, although a formal health economic analysis incorporating device costs, operative time, and overall procedural expenditures will be necessary to determine the net financial impact.

This study had several limitations that should be considered in interpreting the findings. The sample size was modest and may limit detection of smaller effects, particularly at proximal targets. Quantitative assessments of fibrosis, adhesions, and coaptation alignment were not performed. Functional outcomes were inferred from electrophysiological and histological measures rather than behavioral testing. Finally, species-specific differences may limit direct translation to humans.

## Conclusion

Sutureless peripheral nerve repair using Nerve Tape supported durable functional and structural recovery and enhanced distal motor reinnervation compared to conventional neurorrhaphy in a clinically relevant large-animal model. Notably, Nerve Tape resulted in greater electrophysiological signal amplitude at distal targets. These findings support the concept that optimization of the repair interface may improve regenerative yield and overall outcomes following PNI. In summary, Nerve Tape represents a viable alternative to conventional suturing and warrants further evaluation in advanced preclinical and clinical studies.

## Acknowledgments

The authors thank the ULAR veterinary and husbandry staff for animal care. We also thank Jonathan Isaacs and Isaac Clements for the technical assistance.

## Funding

Funding was provided through a Cooperative Research and Development Agreement (CRADA) with BioCircuit Technologies, Inc. Additional support was provided by the Department of Veterans Affairs [Merit Review I01-RX005045 and Center Grant I50-RX004845] and the National Institutes of Health [NIAMS R01-AR083489]. Opinions, interpretations, conclusions and recommendations are those of the author(s) and are not necessarily endorsed by the Department of Veterans Affairs or the National Institutes of Health.

## Conflict of Interest Statement

The authors declare no competing interests.

## Notes

### Competing Interest Statement

The authors have declared no competing interest.

